# GRNUlar: Gene Regulatory Network reconstruction using Unrolled algorithm from Single Cell RNA-Sequencing data

**DOI:** 10.1101/2020.04.23.058149

**Authors:** Harsh Shrivastava, Xiuwei Zhang, Srinivas Aluru, Le Song

**Affiliations:** Department of Computational Science & Engineering, Georgia Institute of Technology, Atlanta, 30332, USA

## Abstract

**Motivation:** Gene regulatory networks (GRNs) are graphs that specify the interactions between transcription factors (TFs) and their target genes. Understanding these interactions is crucial for studying the mechanisms in cell differentiation, growth and development. Computational methods are needed to infer these networks from measured data. Although the availability of single cell RNA-Sequencing (scRNA-Seq) data provides unprecedented scale and resolution of gene-expression data, the inference of GRNs remains a challenge, mainly due to the complexity of the regulatory relationships and the noise in the data.

**Results:** We propose `GRNUlar`, a novel deep learning architecture based on the unrolled algorithms idea for GRN inference from scRNA-Seq data. Like some existing methods which use prior information of which genes are TFs, `GRNUlar` also incorporates this TF information using a sparse multi-task deep learning architecture. We also demonstrate the application of a recently developed unrolled architecture `GLAD` to recover undirected GRNs in the absence of TF information. These unrolled architectures require supervision to train, for which we leverage the existing synthetic data simulators which generate scRNA-Seq data guided by a GRN. We show that unrolled algorithms outperform the state-of-the-art methods on synthetic data as well as real datasets in both the settings of TF information being absent or available.

**Availability:** Github link to `GRNUlar` - https://github.com/Harshs27/GRNUlar

## 1 Introduction

In molecular biology, it is known that the expression level of a gene is controlled by its transcription factors (TFs). Transcription factors are proteins which regulate the expression levels of their target genes in a given cell at a given time. These regulatory relationships can be represented by a graph, called a gene regulatory network (GRN), where nodes represent genes, and an edge from gene A to gene B means that the protein product of gene A is a TF for regulating gene B. This network governs transcription and further decides the way cell behave, and it is of great interest to decipher the interactions in this network.

It has been a long-standing challenge to reconstruct these networks computationally from gene-expression data (Chen *et al.*, 1998; Kim *et al.*, 2003). Recently, single cell RNA-Sequencing (scRNA-Seq) technologies provide unprecedented scale of genome-wide gene-expression data from thousands of single cells, which can lead to the inference of more reliable and detailed regulatory networks (Chen and Mar, 2018; Pratapa *et al.*, 2020). GRNBoost2 (Moerman *et al.*, 2019) and GENIE3 (Vân Anh Huynh-Thu *et al.*, 2010) are among the top performing methods for GRN inference (Chen and Mar, 2018; Pratapa *et al.*, 2020). They are based on fitting a regression function between the expression values of the TFs and other genes. These methods use the information of which genes (corresponding to nodes in the network) are TF genes (thus can have outgoing edges) to achieve better performance. An alternate approach to the inference of regulatory networks can be to pose the sparse graph recovery problem as a graphical lasso problem with *l*_1_ regularization Friedman *et al.* (2008). These approaches are primarily unsupervised in nature and it is not straightforward to include supervised information or prior knowledge of underlying GRNs in their algorithms.

While methods for GRN inference from scRNA-Seq data are improving, simulators which generate synthetic scRNA-Seq data guided by GRNs are also progressing (Dibaeinia and Sinha, 2019; Pratapa *et al.*, 2020). These scRNA-Seq simulators can generate realistic looking data, and have modeled sources of variation in single cell RNA-Seq data, such as noise intrinsic to the process of transcription, extrinsic variation indicative of different cell states, technical variation and measurement noise and bias. The primary application of these realistic simulators is to benchmark the performance of GRN inference methods (Dibaeinia and Sinha, 2019; Chen and Mar, 2018; Pratapa *et al.*, 2020). These evaluations show that the performance of current method for GRN inference is not satisfying even with synthetic data. Indeed, GRN inference is hindered by multiple factors, including the potentially nonlinear relationships between TFs and their target genes, and the intrinsic and technical noise in scRNA-Seq data (Vallejos *et al.*, 2017).

In this paper, we present deep learning frameworks which accommodate the nonlinearity in regulatory relationships, and which are shown to be relatively more robust to technical noise in the data. Moreover, our approach is based on the idea of incorporating prior knowledge of the problem as an inductive bias to design data-driven models (Shrivastava *et al.*, 2018, 2020). We make use of recently proposed unrolled algorithms technique which employs optimization algorithms for the objective function of the problem under consideration as templates for designing deep architectures. The key advantages of unrolled algorithms are (a) few learnable parameters (b) less supervised data points required for training (c) comparable or better performance than existing state-of-the-art methods and (d) more interpretability. Unrolled algorithms have been successfully used in other recent works, including `E2EFold` for RNA secondary structure prediction (Chen *et al.*, 2020) and `GLAD` for sparse graph recovery (Shrivastava *et al.*, 2020). Here we present a novel unrolled algorithm for our deep learning framework, `GRNUlar` (pronounced “granular”, <u>G</u>ene <u>R</u>egulatory <u>N</u>etwork <u>U</u>nrolled <u>a</u>lgorithm), for GRN inference. `GRNUlar` works in the setting where TF information is given. `GLAD` can be applied to GRN inference without using the TF information, and we also provide a modification to it, called `GLAD-TF`, for it to benefit from the TF information.

We utilize the GRN-guided simulators in a novel way: we use a simulator to generate a corpus of simulated data pairs consisting of expression data and the corresponding GRN. This allows us to leverage supervised learning to learn the unrolled algorithm (or neural algorithm) for reconstructing the GRNs from the input gene expression data. It has been argued in the recent works by (Belilovsky *et al.*, 2017; Shrivastava *et al.*, 2020) that a data-driven neural algorithm may be able to leverage this distribution of problem instances, and learn an algorithm which performs better than traditional manually designed algorithms. SERGIO (Dibaeinia and Sinha, 2019) is the most realistic GRN-guided simulator for scRNA-Seq data to date, and is the main simulator we use for training and evaluation of our models. SERGIO generates realistic gene expression data by incorporating known principals of TF-gene regulatory interactions that underlie expression dynamics, and it models the stochastic nature of transcription as well as simulates the non-linear influences of multiple TFs.

We show that our proposed algorithms perform better than state-of-the-art methods on both simulated data and real data from species including human and mouse. Our learned neural algorithms is comparably more robust to high levels of technical noises often observed in realistic settings. We demonstrate that our methods benefit from the supervision obtained through synthetic data simulators, and to the best of our knowledge, we are the first to use the simulators to train neural algorithms for GRN inference from scRNA-seq data.

## 2 Methods

In this section, we first formulate the problem of GRN inference, and briefly introduce some existing approaches. We then present `GRNUlar-base`, which is the “basic” version of `GRNUlar`, and the difference between the two is that the latter uses a technique for initialization and thus improves the runtime of the former. We also provide an insight into a novel loss function which we developed specifically for GRN inference task. Then, we briefly introduce how `GLAD` works for GRN inference and how it is extended to `GLAD-TF`, which makes use of the TF information.

<u>Problem Setting:</u> We consider the input gene expression data to have *D* genes and *M* samples, *X* ∈ *R^M×D^*. Let 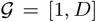 be the set of genes and 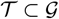 be those which are TFs. We aim to identify the directed interactions of the form (*t, o*), where 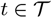 and 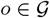.

We note that there can be interactions between TFs themselves. For our method, we assign directed edges between the TFs and other genes and the interactions between TFs are represented by undirected edges. We thus output Completed PDAGs, where PDAGs represent equivalence classes of DAGs and stands for ‘partially directed acyclic graphs’ (Chickering, 2002).

<u>Existing approaches:</u> The common approach followed by many state-of-the-art methods for GRN inference is based on fitting regression functions between the expression values of TFs and the other genes. Usually, a sparsity constraint is also associated with the regression function to identify the top influencing TFs for every gene.

Generally, the form of objective function for GRN recovery used in various methods is a variant of the equation given below

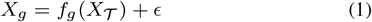

We can view Eq. 1 as fitting a regression between each gene’s expression value as a function of the TFs and some random noise. Simplest model will be to assume that the function *f_g_* is linear. One of the state-of-the-art methods TIGRESS by Haury *et al.* (2012) assumes a linear function of the following form for every gene

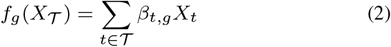

Another top performing method GENIE3 by Vân Anh Huynh-Thu *et al.* (2010), assumes each *f_g_* to be a random forest. GRNBoost2 by Moerman *et al.* (2019) further uses gradient boosting techniques over the GENIE3 architecture to do efficient GRN reconstruction.

### 2.1 GRNUlar: Unrolled model for recovering directed GRNs

We aim to leverage the supervision available to us from the gene expression data simulators. This supervision is in the form of input gene expression data and the corresponding underlying GRN. We speculate that tuning GRN recovery models under this supervision will lead us to better capture the intricacies of the real data and potentially improve upon the unsupervised methods.

We first describe our novel approach of modeling regression functions using a neural network. Based on this modeling, we motivate the use of unrolled algorithms and design its deep architecture.

#### 2.1.1 Modeling regression functions using a Neural Network in a multi-task learning framework

It is intuitive to see that we will be able to better capture dependencies between the expression values in Eq. 1, if we choose *f_g_* to be non-linear function of expression values. Now, it is well known that neural networks (NN) have good capability of representing rich classes of highly non-linear functions. So, we came up with a way of combining the regression formulation with neural networks in a multi-task learning framework (Ruder, 2017), refer Fig. 1.

**Fig. 1:**
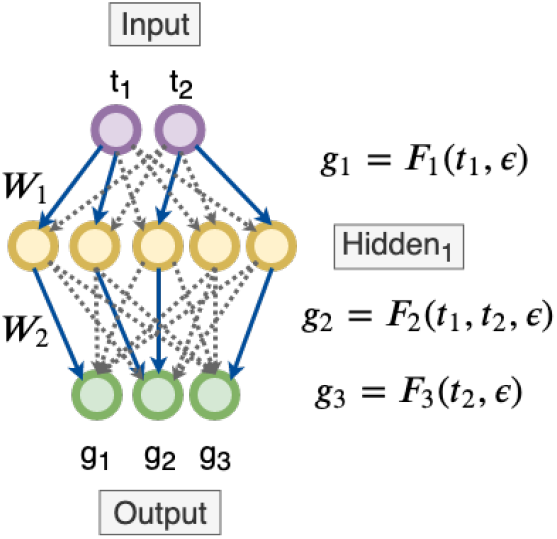
Using the neural network in a multi-task learning framework which act as non-linear regression functions between TFs and other genes. We start with a fully connected NN indicating all genes are dependent on all the input TFs (dotted black lines). Assume that in the process of discovering the underlying sparse GRN our algorithm zeroes out all the edge weights except the blue ones. Now, if there is a path from an input TF to an output gene, then we conclude that the output gene is dependent on the corresponding input TF.

We view the neural networks as a mapping between the input TFs to the output genes. Each output gene is a non-linear function of some combination of input TFs. We view our system as [TF genes → nonL(*W*_1_) → · · · nonL(*W_i_*) → output genes]. If there is a path from an input gene to the output gene, then the output gene is dependent on the corresponding input gene. We thus want sparse NN weights 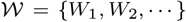 in order to obtain a sparse graph.

We can also easily obtain the dependency matrix between input TFs and output genes as a matrix multiplication, where Θ_*p*_ = Π_*i*_|*W_i_*| represents the matrix product of the neural network weights. This dependency matrix correspond to a GRN. A non-zero value in the matrix Θ_*p*_ indicates that there is an edge between those genes.

This multi-task neural network architecture is an important modeling choice here compared to boosted decision tree based formulations. This model is more expressive than decision trees, and does not need the additional posthoc scoring step. It is also more expressive than a simple non-linear model with additive noise because the NN is jointly optimizing the regression for all the output genes (multi-task learning) and this helps it capture the common dependencies between the TFs and output genes. This also makes the NN model more robust towards external noises.

#### 2.1.2 Motivation for unrolled algorithms

Notice that there can be many neural network representations possible in Section 2.1.1 which can satisfy the Eq. 1 and can lead to different GRNs. These GRNs vary mostly in terms of the sparsity obtained and it is hard for users to manually choose them. Unrolled algorithms help resolve this problem as the sparsity related hyperparameters (eg., the weight of the *l*_1_ norm term) can be learned from supervision.

#### 2.1.3 Designing the unrolled algorithm

We follow similar procedure as the unrolled algorithm designed for the sparse graph recovery task as described in Shrivastava *et al.* (2020). Briefly outlining the approach,

1. Define a related optimization problem for the task at-hand.
2. Apply the Alternative Minimization (AM) algorithm and unroll it for certain number of iterations.
3. Replace the hyperparameters with problem dependent neural networks and treat all the unrolled iterations together as a deep model.
4. Learn this unrolled architecture under supervision by defining a direct optimization objective.

<u>*Step 1:*</u> We consider the following non-linear optimization objective function for the regression defined in Eq. 1 with *l*_1_ penalty

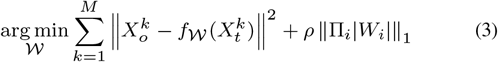

where 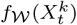 is a neural network. For example, we can define a 2 layer neural network with ‘ReLU’ non-linearity as

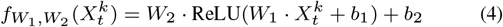

We learn the weights {*W_i_*} and the biases {*b_i_*} while optimizing for Eq. 3.

<u>*Step 2:*</u> We now apply the Alternate-Minimization approach to the optimization given in Eq. 3. Since the objective above is non-linear, we will need an iterative approach to minimize it w.r.t. 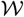. However, we can make our problem easier by separating the two terms such that we can get closed form solution of the *l*_1_ penalty term. We can achieve this by introducing an additional variable *Z* where *Z* = Π_*i*_|*W_i_*|. We then have

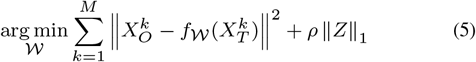

We can include the *Z* term in optimization and as a square penalty term

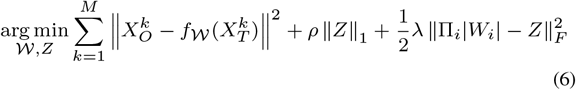

and alternately minimize *Z* and Θ for *l* ∈ [0, *L*] iterations as,

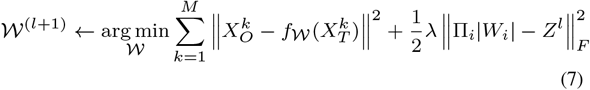

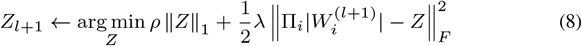

The update of *Z* is of the form *f* (*Z*) + *ρ* ‖*Z*‖_1_, where *f* (*Z*) is a convex function. Similar to Shrivastava *et al.* (2020), the minimizer of this function is the proximal operator given by *η_ρ/λ_*(*θ*) = sign(*θ*) max(|*θ*| − *ρ/λ*, 0).

Thus, the updates of AM algorithm are given by

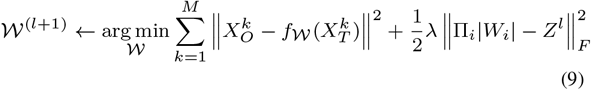

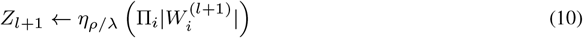

<u>*Step 3:*</u> We identify the proximal operator *η_ρ/λ_* and *λ* as the hyperparameters which control the sparsity of the final graph. We can parameterize them as *ρ_nn_*, *λ_nn_* respectively. These neural networks are minimalist in design and takes the solution of the previous update to predict the next value. We will learn these problem dependent neural network using supervision. As for the Eq. (9), we optimize for the 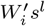 by using standard deep learning optimizers. The corresponding values of *Z^l^* can be obtained by plugging in the 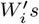 in its closed form update, Eq. 10. We unroll these updates for *L* iterations and treat it as a highly structured deep model.

**Algorithm 1:**
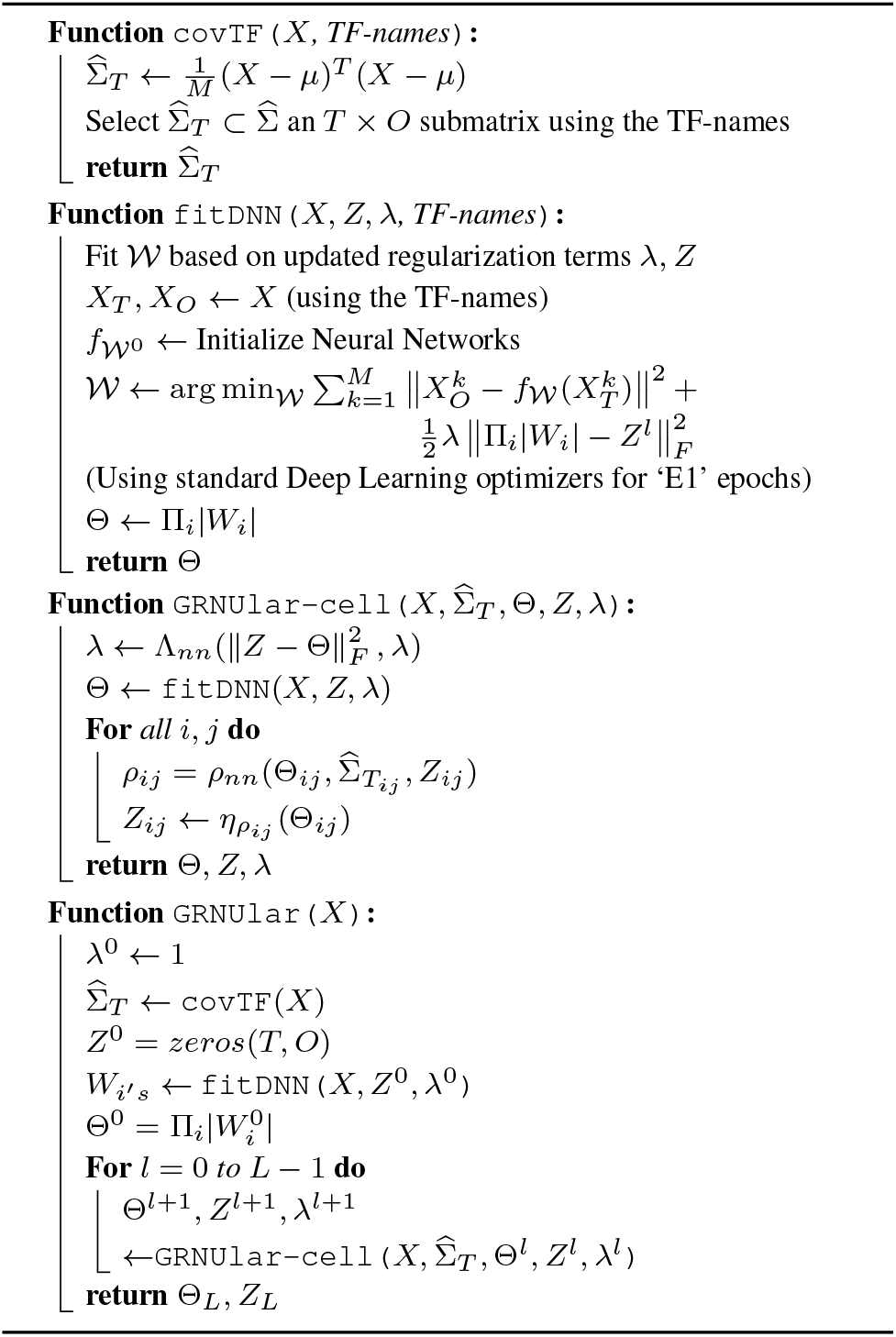
GRNUlar-base

<u>*Step 4:*</u> We optimize the model for minimizing the Frobenius norm and the *F_β_* score between the predicted output graph and the true underlying connection matrix (details of loss function given in Section 2.3).

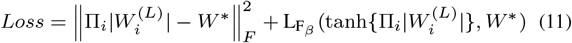

The ground truth *W*\* ∈ {0, 1}^(*O×T*)^ matrix, where 1 indicates an edge between (*t, o*). Algorithm 1: `GRNUlar-base` provides a supervised learning framework for the unrolled model directly based on the updates of the AM algorithm, Eqs. 9 & 10.

A <u>note</u> on backpropagation of gradients: While taking the arg min in the fitDNN function given in Algorithm 1, we consider the *λ* and *Z^l^* as constants. In our PyTorch implementation, we ‘detach’ these variables from the computational graphs while optimizing for 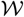. Though, ideally we can retain the computational graph while optimizing but then the memory consumption increases considerably. Another important concern is related to the runtime of Algorithm 1. The `fitDNN` function is called *L* times and it initializes a new neural network each time and minimizes it for the optimization function to a very low error based on the regularization provided by the *λ* and *Z* values. We typically require *E*1 ~ [200, 400] epochs to fit the neural network. This slows down the algorithm significantly. To circumvent this issue, we propose a simple modification to `GRNUlar-base` algorithm described in Section 2.2.

**Algorithm 2:**
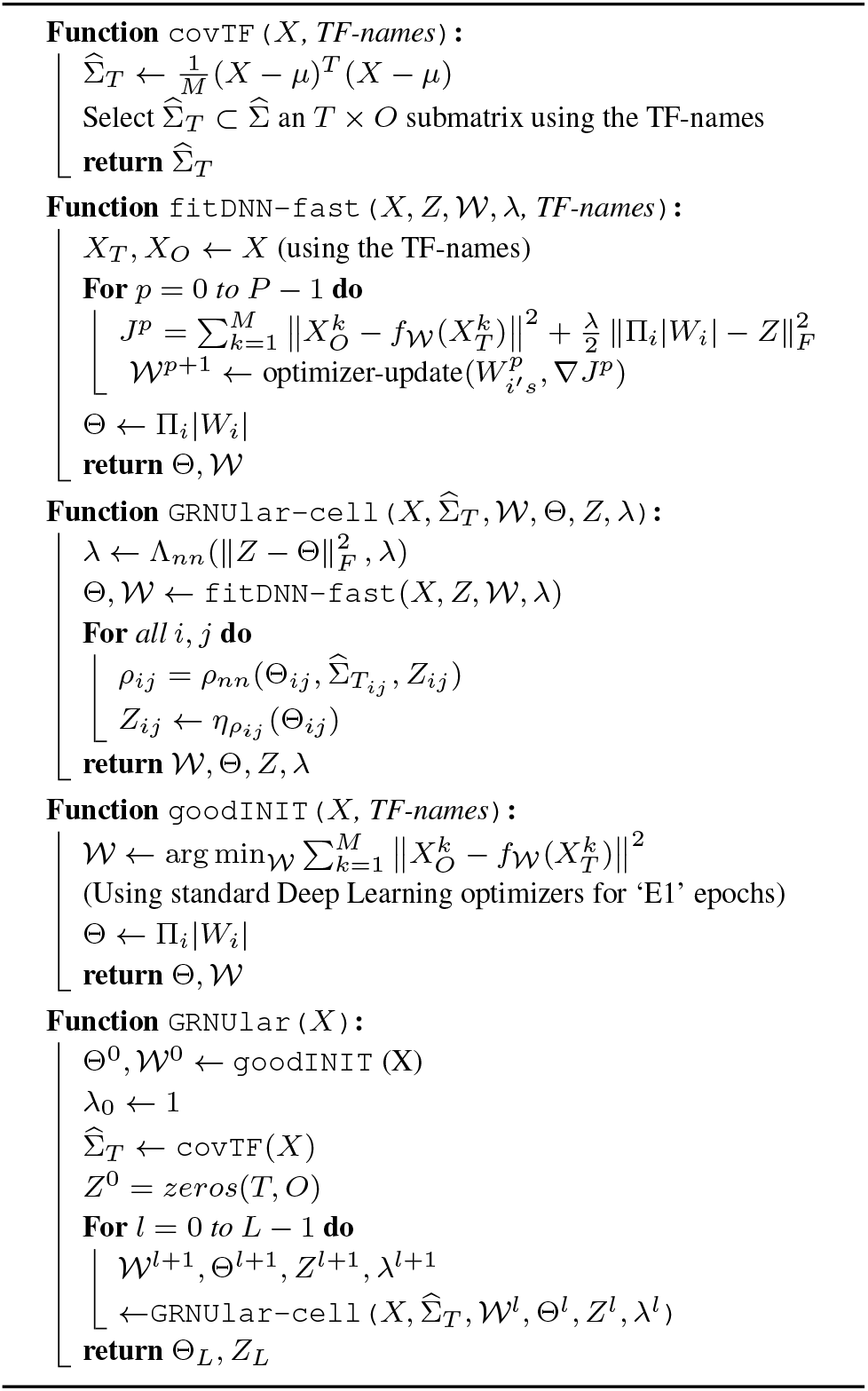
GRNUlar

### 2.2 Efficient GRNUlar algorithm using ‘good’ initialization

We propose an alternate ‘good’ initialization technique to reduce the runtime of Algorithm 1. We posit that, if we optimize for the 1^*st*^ term of Eq. 9 beforehand and obtain good initial values of 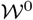, we then just need to do minor adjustments to the 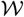 as we update *Z* and *λ*. We then just need to unroll the optimization (the new `fitDNN-fast` function) for only few iterations *P* ~ {2, 5, 10}. Refer Algorithm2 as well as Fig. 2 which pictorially shows the highly structured deep architecture of the `GRNUlar` algorithm. For the *p^th^* unrolled iteration, we have

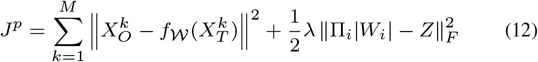

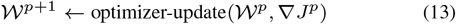

**Fig. 2:**
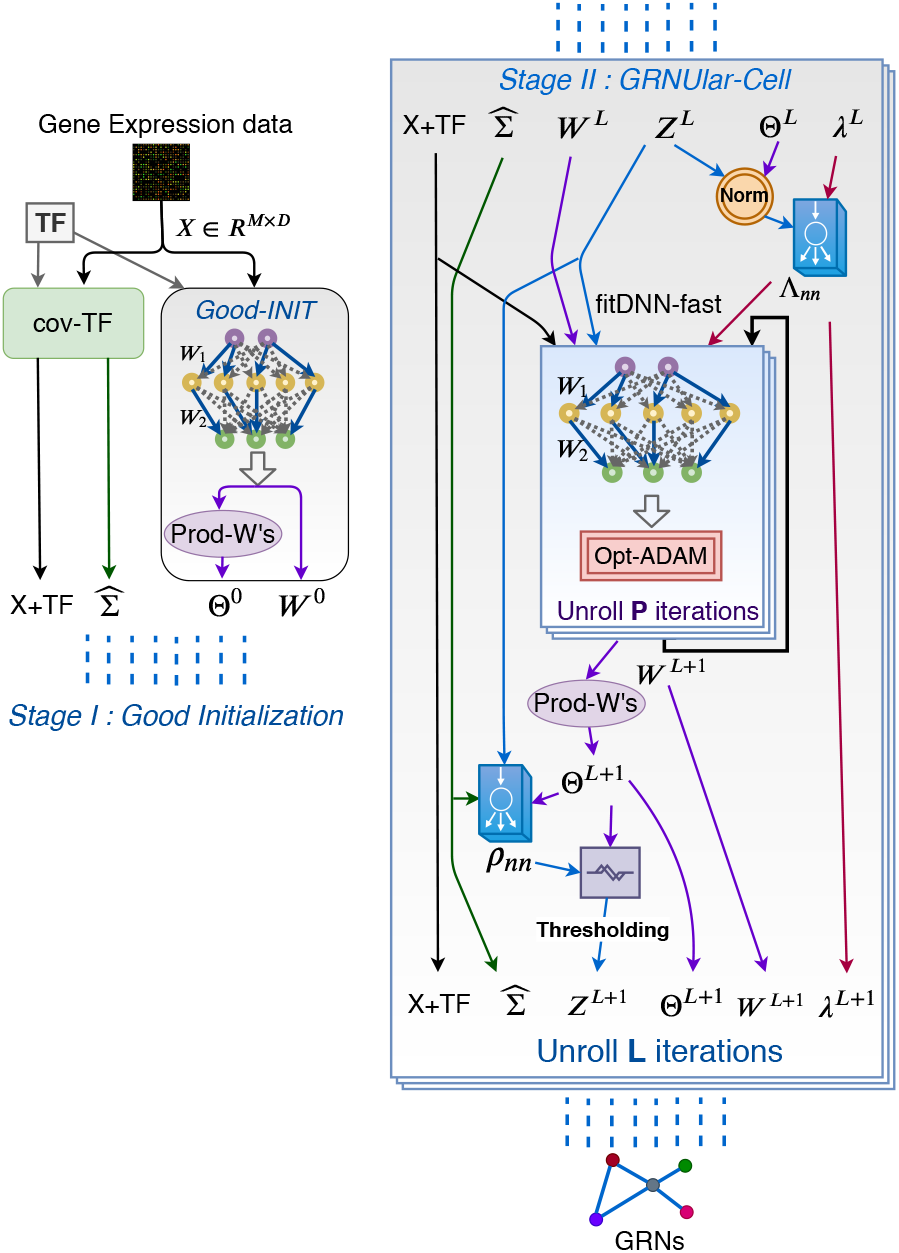
Visualizing the `GRNUlar` algorithm’s architecture.

This is a significant reduction in the number of unrolled iterations required over the `fitDNN` function. The `GRNUlar-cell` in Algorithm 2 also does not require large number of unrolled iterations *L*. The neural network *ρ_nn_* is learning the entrywise thresholding operation and *λ_nn_* learns to update its value from norm difference and its previous value. We observe that in every iteration of Algorithm 2 we optimize the unrolled parameters *ρ_nn_*, *λ_nn_* (tiny neural networks) to learn the underlying graph sparsity from the supervision provided. Thus, we want to <u>highlight</u> that the overall training does not require much training data as well as the number of unrolled iterations. We also empirically verify that `GRNUlar` algorithm performs equivalent to `GRNUlar-base` algorithm with significant runtime improvement.

A <u>note</u> on the general idea of neural network based parameterization: For the `GRNUlar` algorithm, we can further parameterize the optimizer update given in Eq. 13 and learn it from the supervision provided, similar to Λ_*nn*_. In our current implementation, we use the ‘adam’ optimizer. We want to highlight that our technique of parameterization in an unrolled fashion is very generic and can be used for any off-the-shelf optimizer. For instance, consider the example of parameterizing gradient descent optimizer which is realized using the Λ_*nn*_ update. We just need to define the neural network based parameterization in a way that is more generic then the optimizer’s update equation. The neural network based update 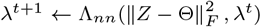 subsumes the standard gradient descent update given by 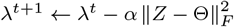.

### 2.3 Training GRNUlar

The GRN graphs are usually quite sparse and so we want our loss function to be robust enough to recover sparse edges. Since, there are multiple metrics like precision, recall, F1 score etc. which are commonly used for evaluation purposes of the recovered graphs, it will be very useful if we can define a loss function which can find a desirable balance between them.

To address the above concerns, we develop a differentiable version of the *F_β_* score. Say, Θ^*p*^ represents our predicted graph (adjacency matrix) and the true underlying graph is represented by Θ* and we assume that all the entries of Θ ∈ [0, 1]. So, we can write the true positives(TP), true negatives(TN), false positives(FP) and false negatives(FN) as follows:

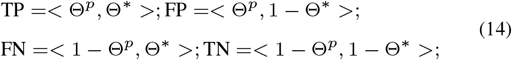

where < ·, · > represents matrix inner product, which is the summation of entry-wise products. Based on the above differentiable representations, we define differentiable *F_β_* score and the corresponding loss function as:

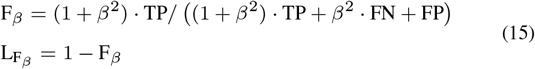

Eq. 15 tells us that having *β* > 1 will weigh recall higher than precision as it places more emphasis on the FNs. Similarly, having *β* < 1 will attenuate the influence of FNs and thus weigh recall lower than precision.

A <u>note</u> on inputs to loss function: In order to ensure that the entries of Θ^*p*^ ∈ [0, 1] we pass it through the 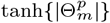 operation which is an entry-wise absolute value followed by an entry-wise tanh function. In some cases, we also do an additional diagonal masking operation, 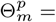 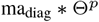, as we want to ignore the self loops.

Thus, we define a loss function between the predicted Θ^*p*^ and true underlying matrix Θ* as the combination of the MSE (or Frobenius norm) loss and the 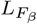 loss. It is often tricky to jointly optimize and balance between multiple loss functions. Taking hint from the loss balancing technique described in Rajbhandari *et al.* (2019), we introduce a balancing ratio 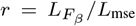 which adjust the scales of both the losses. Note that while implementing this scaling, at every epoch, we calculate ‘r’ by detaching the losses from the computational graph to keep it as a constant.

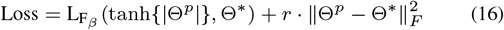

For training `GRNUlar`, a collection of input expression data and their corresponding ground truth GRNs can be sampled from GRN guided realistic data simulators. The loss function in Eq. 16 is designed to directly optimize the output GRN prediction of `GRNUlar` to the ground truth GRN connections. We aim to optimize the loss function over the average of such data pairs so that the learned architecture is able to perform well over a family of problem instances.

### 2.4 GLAD model & proposed modification for TFs

The `GRNUlar` algorithm described above is designed to reconstruct GRNs using the TF information. But, there can be cases where we do not know the underlying TFs. Majority of the existing methods are typically designed to include TF information and their performance in terms of recovery quality and runtimes deteriorate significantly if the TFs are not provided. Our experiments corroborate these statements.

We found an alternative method to slightly mitigate above concerns. We directly use the unrolled model `GLAD` (Shrivastava *et al.*, 2020) for the GRN inference problem. This model does not use the TF information for GRN recovery. Briefly, the `GLAD` model’s architecture is based on the unrolling the iterations of an Alternate Minimization algorithm for the graphical lasso problem with applications towards sparse graph recovery. We refer the reader to the Algorithm 3 in Supplementary-A

<u>`GLAD-TF` model:</u> We modify the architecture of `GLAD` to include prior information about the TFs if available. We assume that all the edges in the actual GRN have at least one gene which belongs to the list of TFs. We introduce a masking operation at every step of the unrolled iterations as shown in Algorithm 4 in Supplementary-A, which eliminates all the unwanted edges that are between the non-TF nodes.

## 3 Experiments

### 3.1 Description of evaluation metrics & methods

<u>Recovery Metrics:</u> Following the metrics used in (Dibaeinia and Sinha, 2019; Chen and Mar, 2018), we also use AUROC (Area Under the Receiver Operating Characteristics) and AUPRC (Area Under the Precision Recall Curve) values for our evaluation.

<u>Baseline methods:</u> We compared with the GRNBOOST2, GENIE3 as they are representative of regression based methods. We used the Arboreto package to run these algorithms (Moerman *et al.*, 2019). We additionally compared with the Graphical Lasso algorithm (GLASSO) using their “scipy” implementation. We did extensive fine tuning of the hyperparameters for all the baseline methods using the training/valid data and then reported the results on the test data.

<u>Settings of the unrolled algorithms:</u> For the `GLAD` model, we used the standard initialization as recommended by the authors (refer Fig.3). We chose the number of unrolled iterations *L* = {15, 30}. For the `GRNUlar` model, we used the standard initialization of the thresholding parameters *ρ_nn_*, *λ_nn_*. Now, we need to decide the dimensions of neural network (NN) which fits the regression between the input TFs and the output genes in the `fitDNN-fast` function. Our general strategy is to have number of hidden layers, *depth ≥* 2, of the NN. We roughly choose the number of hidden units in layer ‘j’ as *H_j_* ≥ 4 · *H*_*j*−1_ and we also satisfy #TF ≤ *H*_1_. The number of epochs to fit the regression is decided using the results on the validation dataset. We empirically observed that we need around [200, 500] iterations to fit the neural network in the `goodINIT` function. We chose the unroll parameters *L* = 15 and the values of *P* = {2, 5, 10, 20} with 2 or 3 layers neural network for our experiments.

**Fig. 3:**
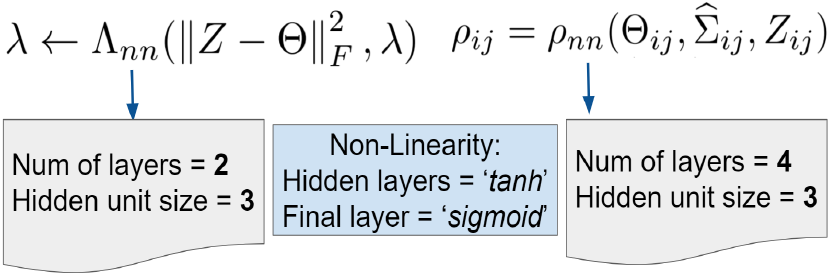
**Minimalist** neural network architectures designed for the problem dependent parameterizations *ρ_nn_* & Λ_*nn*_ of both the unrolled models (taken from `GLAD` work).

For both the unrolled methods, we chose 2 models based on AUPRC and AUROC results on the validation data. We use the scaled loss function (Eq. 16) to jointly optimize for MSE and *F_β_* loss. The values of *β* used in our experiments were chosen from the set {0.5, 1, 2, 5}. We implemented the unrolled algorithms using PyTorch and were ran on Nvidia P100 GPUs.

### 3.2 Evaluating GRN inference methods on synthetic data

In this subsection, we aim to conduct an exploratory study to gauge the generalization ability of the unrolled algorithms for the GRN inference task using the synthetic data simulator. We compare all the algorithms on various simulator settings and list our key observations.

<u>*Graphs and expression data for training:*</u> To provide supervision for the unrolled algorithms we use the SERGIO simulator. To create random directed graphs (GRNs) we first decide on some number of TFs or master regulators. Then, we randomly add edges between the TF and the other nodes based on the sparsity requirements. Also, we randomly add some edges between the TF themselves but exclude the self-regulation edges. We further add minimal number of edges to maintain connectivity of graph as it is a requirement for the simulator. This graph is then given as input to the SERGIO simulator to generate corresponding gene expression data. For the experiments in this subsection, we take train/valid/test=20/20/50 graphs respectively with number of genes D=100. All these graphs were sampled from similar settings. We usually choose the ratio of TF with total number of nodes=0.1 (~ 10 TFs for D=100) and sparsity of training graphs=0.1. We want to highlight that we do not need many graphs to train the unrolled models as we are primarily learning the sparsity pattern from supervision and need small neural networks for the same, refer Fig.3.

From literature on sample complexity theory of sparse graph recovery, eg. (Ravikumar *et al.*, 2011), we know that recovery of the underlying graph improves with the increasing number of samples. Hence, we ran our experiments with varying number of the total single cells, M=[100, 500, 1K, 5K, 10K]. We also observed that varying the number of cell types (corresponding to the number of clusters of the cells) of the SERGIO simulator considerably affects the GRN inference results, so we also evaluated the methods by varying the number of cell types of the simulator C=[2, 5, 10]. Note, we adjusted the number of cells per cell type to maintain same total number of cells.

<u>*SERGIO parameter details:*</u> SERGIO provides a list of parameters to simulate cells from different types of biological processes and gene-expression levels with various amount of intrinsic and technical noise. We simulate cells from multiple steady states. When simulating data with no technical noise (what we refer to as clean data), we set the following parameters: *sampling-state*=15 (determines the number of steps of simulations for each steady state); *noise-param*=0.1 (controls the amount of intrinsic noise); *noise-type*=“dpd" (the type of intrinsic noise is Dual Production Decay noise, which is the most complex out of all types provided); We set genes’ decay parameter to 1. The parameters required to decide the master regulators’ basal production cell rate for all cell types - low expression range of production cell rate ~ *U* [0.2, 0.5] and high expression range of cell rate ~ *U* [0.7, 1]. We chose *K* ~ *U* [1, 5], where ‘*K*’ denotes the maximum interaction strength between master regulators and target genes. Positive strength values indicate activating interactions and negative indicates repressive interactions and ±1 signs are randomly assigned. When adding technical noise, we add the dropout events which are considered to be a major source of technical noise in real data. Parameters which control the amount of dropouts include *shape* (which we set to 20) and *percentile*, which we vary among the values *q* = {25, 50, 75}. Larger *q* corresponds to higher technical noise. All other parameters are set to default values.

For experiments in this subsection, we found that `GRNBoost2` consistently outperforms `GENIE3` method and hence we only show `GRNBoost2` results in the plots. Also, each data point in the plots represent its value along with the standard deviation over the test graphs.

#### 3.2.1 Clean: simulated data with no technical noise

The ‘clean’ gene expression data from SERGIO follows all the underlying kinetic equations but excludes all the external technical noises. We can consider this data as being recorded with no technical errors. Fig. 4 compares different methods on their AUPRC performance on varying number of cells and number of cell types. For `GRNUlar` model we chose 2 layer NN with P=5, *H_d_*={40, 60, 100}, L=15, vary *β* = {2, 5} in the loss function and we selected the best model based on the validation results.

**Fig. 4:**
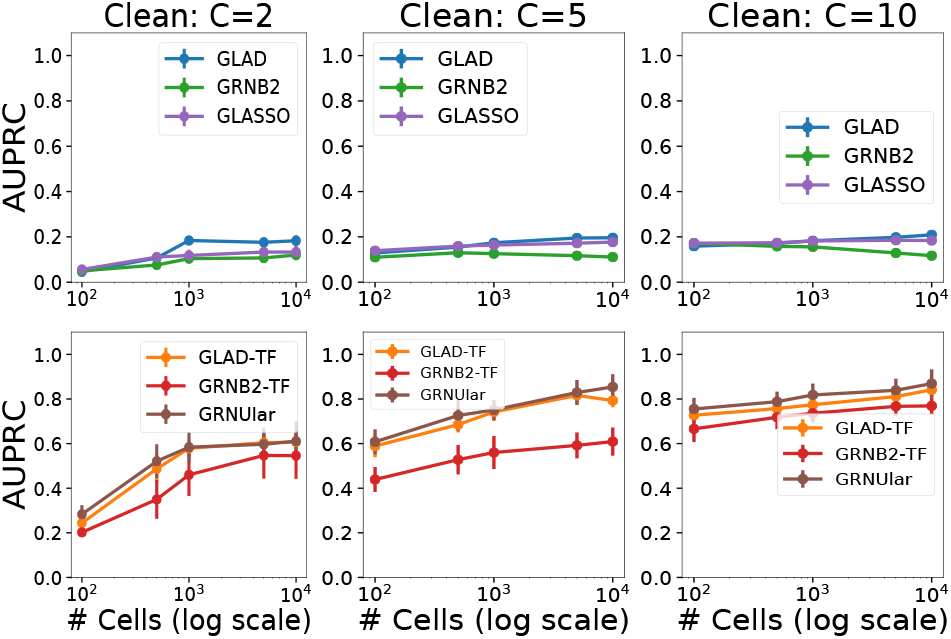
Clean data setting of the SERGIO simulator with D=100 genes. As the number of cell types increase from C=2 to C=10, we see that the AUPRC values increase in general. The top panel contains the methods without the TF information and the bottom panel shows the AUPRC plots of the methods including the TF information. The unrolled algorithms in general outperform the traditional methods.

<u>Key highlights:</u> (a) We observe that `GLAD` works better than `GRNBoost2` in settings where TFs are absent. (b) The unrolled algorithms consistently outperform others in no-noise settings, with and without TFs.

#### 3.2.2 Noisy: simulated data with technical noise

We evaluate the performance of unrolled algorithms on the more challenging and realistic noisy settings. We limit varying the technical noise to dropouts while keeping the default settings for other SERGIO parameters. For higher levels of dropouts, researchers sometimes resort to data imputation techniques (which attempt to recover the number of molecules being dropped) as a preprocessing step which marginally improves results. For these experiments, we report the results without the imputation preprocessing step and compare all methods directly on the noisy data obtained from the simulator. We also note, that while training the models, we train on data with low dropout rates *q* = {0, 25} and use the same models to predict networks on data with higher dropout rates.

Fig. 5 shows the performance of the methods without the TF information. We can see that as the dropout percentile increases, all the methods start to fail significantly. For example, the system becomes too ill-conditioned for the graphical lasso method.

**Fig. 5:**
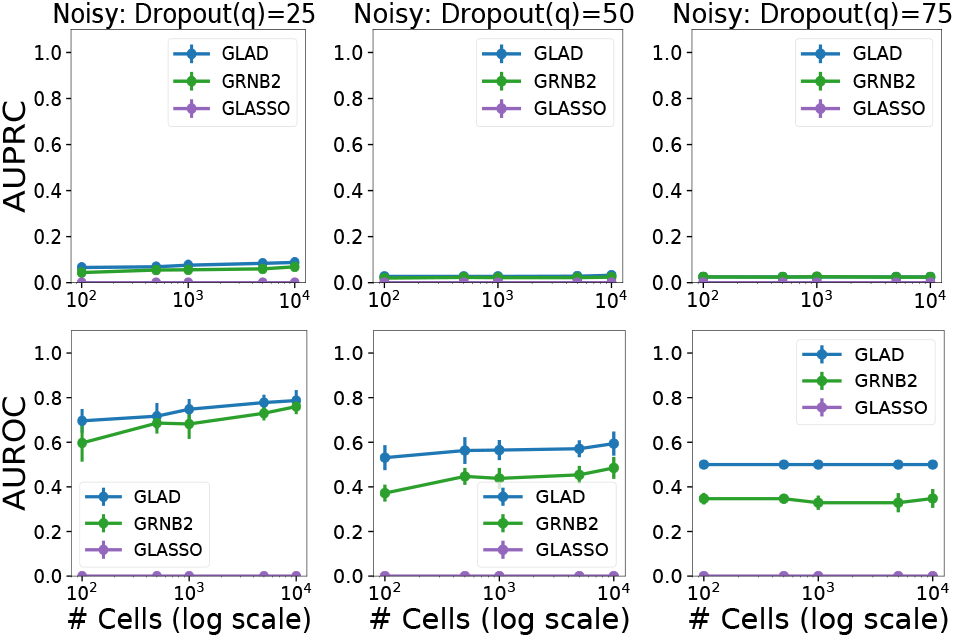
(Without TF information) : Noisy data setting with D=100 and C=5. We vary the dropout percentile values as [25, 50, 75] in both the upper panels (AUPRC values) and the lower panel (AUROC values). Again, in the case of no TF information provided, `GLAD` outperforms other methods.

Fig. 6 includes the TFs. As we move towards the right, dropout percentile increases and we can observe the deterioration in AUPRC values. Although, the `GRNUlar` model’s AUROC performance is quite robust with increasing dropouts.

**Fig. 6:**
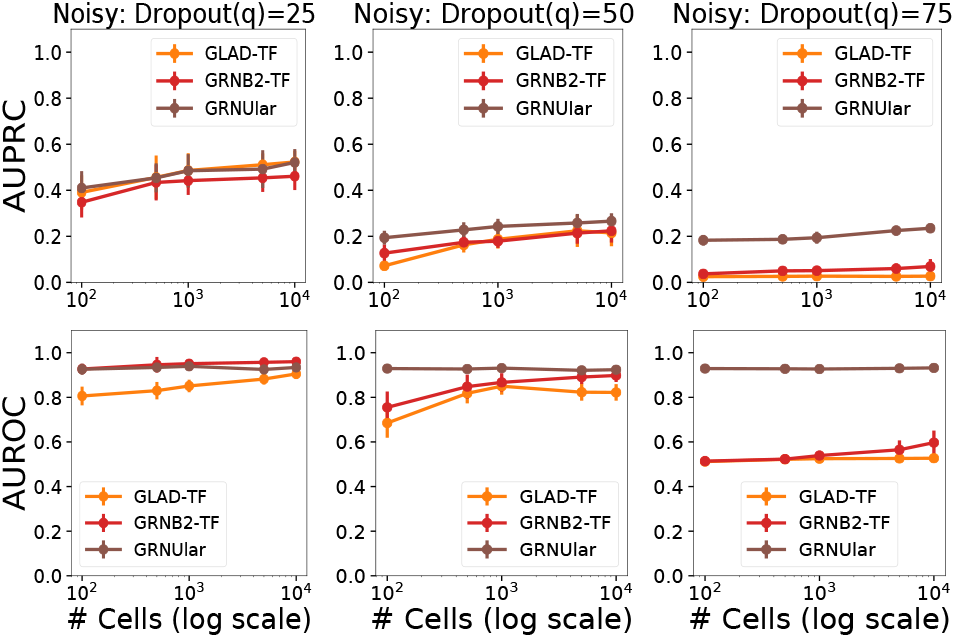
(With TF) : Noisy data setting of the SERGIO simulator with dropout-shape=20, D=100 and C=5. We vary the dropout percentile values as [25, 50, 75] in both the upper panels (AUPRC values) and the lower panel (AUROC values). `GRNUlar` has a clear advantage in noisy settings. We also observe that the performance of `GLAD-TF` is close to the GRNBoost2 method in terms of AUPRC but falls a little behind in AUROC metric.

<u>Key highlights:</u> (a) The case where we do not provide the TF information, then the `GLAD` method outperform others. (b) For the case with TF information, the new unrolled algorithm `GRNUlar` significantly outperforms its competitors. Even in the case of 75% dropout value, where the other algorithms almost give output equivalent to random prediction, `GRNUlar` is able to handle this high percentage of missing information.

### 3.3 Realistic data from SERGIO: Ecoli & Yeast

The challenge for the data-driven models is to be able to generalize to real datasets. Thus, it is important for us to test the ability of the unrolled algorithms to generalize over different settings from that of the training.

To perform this study, we make use of the realistic data sets provided by the SERGIO simulator. They provide three scRNA-Seq datasets DS1, DS2 & DS3 which are generated from input GRNs with 100, 400 & 1200 genes respectively. These networks were sampled from real regulatory networks of *E.coli* and *S.cerevisae*. For each dataset, the settings are: number of cell types C=9; total number of single cells M=2700, and there are 300 cells per cell type. Each data set was synthesized in 15 replicates by re-executing SERGIO with identical parameters multiple times. The parameters were configured such that the statistical properties of these synthetic dataset are comparable to the mouse brain in (Zeisel *et al.*, 2015).

We define the our training and testing settings such that there are considerable differences between the training and testing datasets. We use all of the DS1, DS2 & DS3 datasets for testing, and only the DS1 dataset for training. The major similarities and differences between the training and testing data are:

<u>Similarities</u>: (1) The SERGIO settings for the train data are sampled from similar ranges as that of DS1. Specifically, the parameters like production cell rates, decays, noise-param and interaction strength. (2) We maintain similar number of genes as in test graphs for training.
<u>Differences</u>: (1) The underlying GRNs are completely different in terms of sparsity and connection patterns. (2) We train on data with no dropouts as opposed to 82% dropout percentile in the case of the DS datasets. We thus, keep the dropout information hidden from the models. (3) The datasets DS2, DS3 are completely different from train data (& DS1) in terms of the underlying GRN as well as the corresponding SERGIO parameters are sampled from different range of values. For details, refer to Table1 in Dibaeinia and Sinha (2019) & supplementary Tables S1, S3.

`GRNUlar` settings: We used the parameter settings as mentioned in Section 3.1. We used a 2 layer neural network in the `fitDNN-fast` function for these experiments with a single hidden layer *H*_1_. Following our strategy to choose the dimensions, *H*_1_ ~ 4 × #TF. The number of TFs in DS1/DS2/DS3 are 10/37/127 respectively. So, we chose *H*_1_ ~ 40/200/500 respectively as the hidden layer dimensions.

Fig. 7 re-highlights that the key observations in the previous sections holds true for these set of experiments as well. While there is significant amount of differences between the training and testing data, `GRNUlar` in general performs better than other methods in the noisy settings. We can improve its performance further, by training using multiple data simulators and by better finetuning of simulators to resemble the real data.

**Fig. 7:**
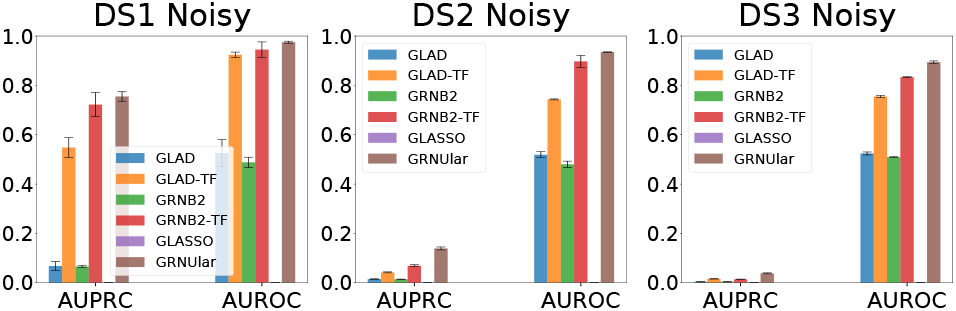
(Noisy settings, dropout percentile=82%) We report the average results over 15 test graphs in the noisy settings. `GLAD` is the best performing method in absence of TFs. `GRNUlar` gives notable AUPRC values and it outperforms other methods.

### 3.4 Real single cell RNA-Seq datasets

In this section, we evaluate the methods on seven datasets from five experiments which include human mature hepatocytes (hHEP) (Camp *et al.*, 2017), human embryonic stem cells (hESC) (Chu *et al.*, 2016), mouse embryonic stem cells (mESC) (Hayashi *et al.*, 2018), mouse dendritic cells (mDC) (Shalek *et al.*, 2014), and three lineages of mouse hematopoietic stem cells (Nestorowa *et al.*, 2016): erythroid lineage (mHSC-E), granulocyte-macrophage lineage (mHSC-GM) and lymphoid lineage (mHSC-L). These are the same datasets used in Pratapa *et al.* (2020) and we use their corresponding ground-truth networks for our experiments as well. For each dataset there are three versions of ground-truth networks: cell-type-specific ChIP–seq, nonspecific ChIP–seq and functional interaction networks collected from STRING. We then have in all 21 different data pairs, 7 different types of expression data evaluated against 3 different types of ground truth.

<u>Preprocessing:</u> For each gene expression data and its corresponding network, we first sort all the genes according to their variance and select the top 500 varying genes. We also have access to a list of known TFs. We only consider all the TFs whose variance had p-value at most 0.01. Now, we find the intersection between the top 500 varying genes and all the TFs to find the subset of genes which act as the TF, see Table 3 in Supplementary-B. Then, we select the sub-graph of top 500 varying genes from the underlying GRN as our ground truth for evaluation.

<u>Training details:</u> We train on the expression data which is similar to the SERGIO settings for the DS2 dataset as it has similar number of genes as the real data. We chose the underlying GRNs for supervision as the random graphs described in Section 3.2. We fixed the number of genes D=400, the number of cell types C=9 and total number of single cells M=2700. The `GRNUlar` settings were *P* = {2, 5, 10}, *L* = 15, *H_d_* = {200} with 2 layers neural network.

Fig. 8 shows the heat map with of the AUPRC and AUROC values for the 21 data pairs. In general for real data, we observe very low AUPRC values, this is primarily due to the highly skewed ratio between true edges and total possible edges (Chen and Mar, 2018; Davis and Goadrich, 2006). We can observe that the unrolled algorithm `GLAD-TF` gives slightly improved performance over `GRNBoost2` and `GENIE3` methods in terms of AUPRC. the `GRNUlar` algorithm clearly outperforms other methods in all test settings. We can further improve the results by tuning the SERGIO simulator settings closer to the real data under consideration.

**Fig. 8:**
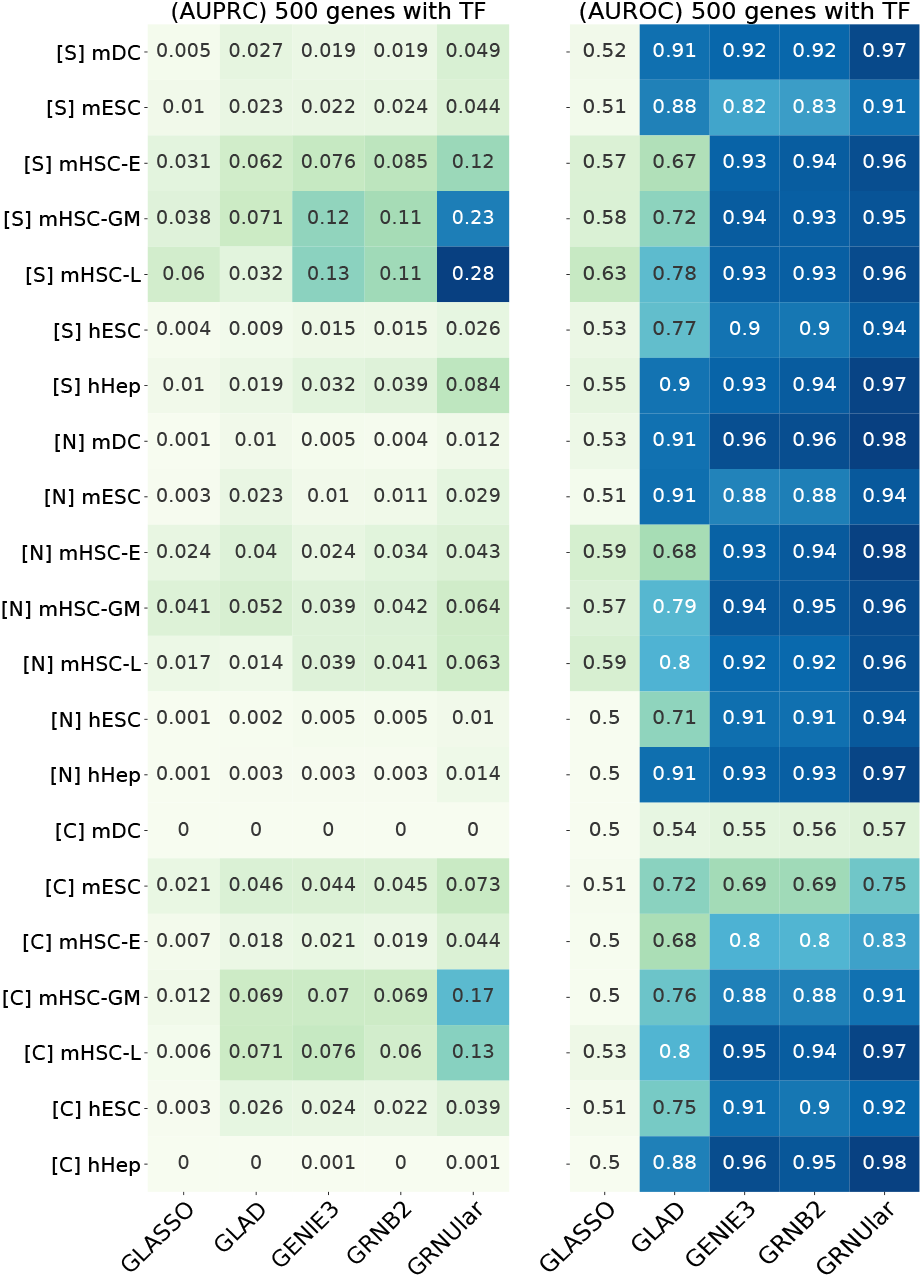
Heatmap of AUPRC and AUROC values of the real data from the BEELINE framework by Pratapa *et al.* (2020). We ran all the methods including the TF information. [S]/[N]/[C] represent the ground truth networks [String-network]/[Non-Specific-ChIP-seq-network]/[Cell-type-specific-ChIP-seq] respectively. Data of the species [m] mouse and [h] human were used. `GRNUlar` performs better than the other algorithms in both the metrics.

We analysed the network predicted by `GRNUlar` from the mESC dataset. We chose TFs and genes corresponding to gene ontology (GO) terms related to ESC cell differentiation and cell fate towards endodermal cells as in this dataset the ESC cells are induced to differentiate into primitive endoderm cells (Hayashi *et al.*, 2018). From BioMart (Kinsella *et al.*, 2011) we obtained 286 genes. We took the intersection between these genes with our predicted GRN (with 500 genes) and got 32 genes.

We first compared the interaction scores predicted by `GRNUlar` among all 500 genes and the scores among the 32 genes, without applying any threshold. We found that the latter set of scores is significantly higher than the former set of scores (Fig. 9). This means that there are more regulatory activities among the genes related to the expected biological processes compared to all the genes selected by variation. We then set the threshold for interaction score as 0.22, and obtained the network shown in Fig. 11. In this network, the TFs SOX7, SOX17, MTF2, GATA6 and CITED2 are known TFs in either stem cell differentiation or embryo development; NOTCH1 and RBPJ are TFs in the NOTCH pathway which controls cell fate specification (www.genecards.org). The TFs with highest interaction scores are highly relevant TFs for the cells under study.

**Fig. 9:**
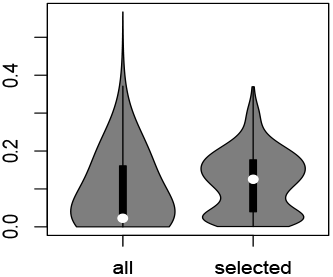
Violin plot comparing the scores of all interactions in the 500 genes (left) and scores of interactions between the 32 genes (right). Wilcoxon p-value is 1.3e-14.

**Fig. 10:**
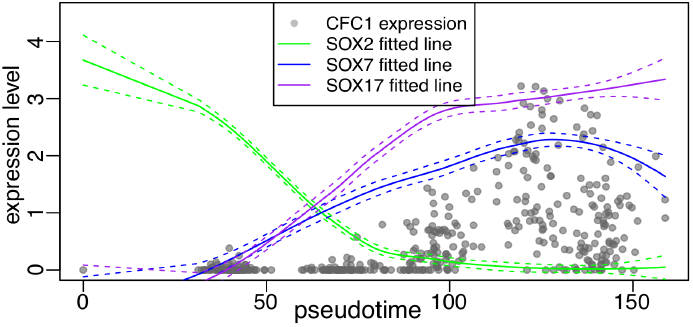
Comparison of gene-expression patterns over the pseudotime for CFC1 and the SOX family TFs.

**Fig. 11:**
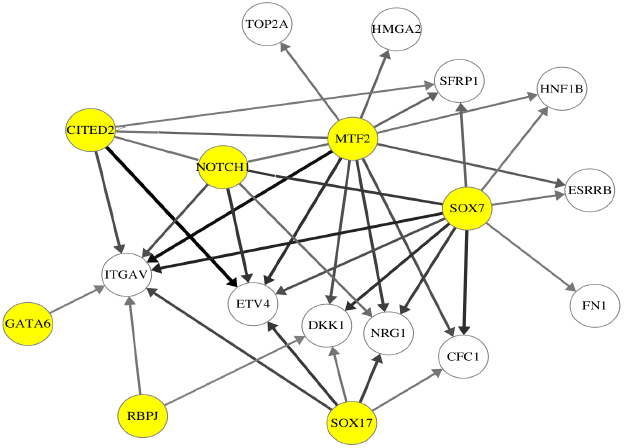
A subnetwork (CPDAG) with genes related to stem cell differentiation from `GRNUlar` predicted network. TFs are the nodes with yellow background. Darker edges mean higher predicted score for the interaction.

We now show how our predicted interactions may bring new biological insights. For instance, we noticed that one of the target genes of SOX7 with strong interaction is CFC1. From ChIP-Seq experiments (the [Cell-type-specific-ChIP-seq] ground truth network mentioned previously), SOX2 is a TF for CFC1. However, in our prediction results, we predicted SOX7 and SOX17 as the TFs for CFC1. We note that the dataset consists of ESC cells differentiating into primitive endoderm cells, and SOX2 is a key TF in mouse ESCs governing the pluripotency of the cells (Masui *et al.*, 2007). As the cells differentiate, the pluripotency goes down, so the SOX2 function may also decrease. To verify this, we use the pseudotime of the cells obtained from (Pratapa *et al.*, 2020), which was inferred with Slingshot (Street *et al.*, 2018), and visualize the gene-expression levels of CFC1, SOX2, SOX7 and SOX17 (Fig. 10). For readability we plot the actual gene-expression levels cell by cell only for CFC1, and for the SOX TFs we plot the fitted lines of their expression levels obtained using LOESS regression. The dashed lines represent the standard deviation. We see that indeed the SOX2 expression decreases along the pseudotime, and the expression levels of CFC1, SOX7 and SOX17 increase. The fitted lines of SOX7 and SOX17 show that they are much better predictors for the expression of CFC1 than SOX2. Indeed, it is discussed that SOX7 and SOX17 are highly related members of the SOX family and their high expression in ESCs are correlated with a downregulation of the pluripotency and an upregulation of the primitive endoderm-associated program (Sarkar and Hochedlinger, 2013). This example showcases how we can use predicted regulatory networks to find regulatory pathways for a specific biological program. Some of these may already have evidence in literature but some may be new and our prediction can be used to provide hypothesis for further experimental validation.

### 3.5 Runtimes of different methods

Tables 1 & 2 show the inference time required for different methods with the TF information included. We run different methods on different platforms and hence comparing them directly is not fair. Although, we include them to give an idea of the runtimes to the reader. The traditional methods takes multiple hours to run in absence the TF information.

**Table 1.** Inference runtimes for the `GRNUlar` model with 2 layer NN, as we vary the hidden layer dimensions *H_d_*. The time is reported for one complete forward call (`goodINIT` and `fitDNN-fast`) for D=1200 genes graph. Other relevant parameters settings were P=5, L=15, DNN epochs E1=400.

| Time (secs) | $H_d=200$ | $H_d=500$ | $H_d=1000$ |
| --- | --- | --- | --- |
| GRNular [gpu] | 0.89 | 1.33 | 2.10 |

**Table 2.** Inference times for different methods on D=1200 genes graph. The unrolled algorithms were ran on GPUs (NVIDIA P100s) while the traditional methods were ran on CPU having a single node with 28 cores.

| Methods | GLASSO [cpu] | GRNB2 [cpu] | GENIE3 [cpu] | GLAD [gpu] | GRNular [gpu] |
| --- | --- | --- | --- | --- | --- |
| Time (secs) | 180 | 612 | 1020 | 0.79 | 1.33 |

## 4 Conclusions and Discussions

We present a novel unrolled algorithm `GRNUlar`, for the inference of gene regulatory networks from scRNA-Seq data. The `GRNUlar` model’s deep architecture takes its inductive bias from the regression based formulation of the GRN recovery problem. We make use of a neural network in a multi-task learning framework to model the regression between TF and other genes. In cases where the TF information is available, we show that `GRNUlar` consistently performs better than representative existing methods under various settings of both simulated data and real experimental data, especially in the more realistic case of added technical noise. The deep learning framework accommodates the nonlinearity of the regulatory relationships and provides tolerance to the technical noise in the data. We also show the superior performance of a recently developed unrolled algorithm `GLAD` in absence of TFs.

The methods we propose are the first supervised deep learning methods for GRN inference from scRNA-Seq data. Our models learn the charateristics of the underlying GRNs through the simulated data from GRN-guided simulators like SERGIO, and demonstrate the application of these simulators in training deep learning models apart from benchmarking computational methods. Similar techniques can be investigated, not only for the task of GRN inference, but also for other analysis tasks for scRNA-Seq data like clustering and trajectory inference (Luecken and Theis, 2019), by using the available realistic simulators for scRNA-Seq data (Zhang *et al.*, 2019; Dibaeinia and Sinha, 2019).

## Acknowledgements

We thank Aditya Pratapa and Prof. T. M. Murali for sharing the gold standard networks for real data used in their paper Pratapa *et al.* (2020).

## Supplementary Material

### A: GLAD and proposed modification

The `GLAD` work formulates the sparse graph recovery problem (undirected graphical model) for the GRN recovery as fitting a multivariate Gaussian distribution on the input gene expression data with a *L*1 normalization term. It is based on the unrolling the iterations of an Alternate Minimization algorithm for the graphical lasso problem with applications towards sparse graph recovery, refer Algorithm 3.

**Algorithm 3:**
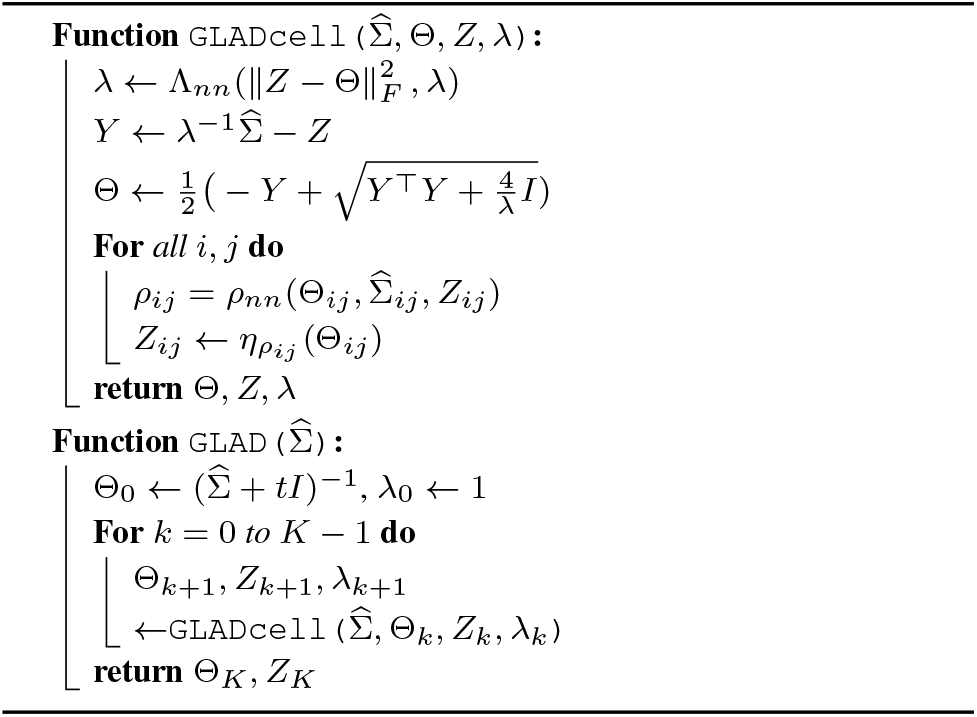
GLAD

<u>`GLAD-TF` model</u>: We modify the architecture of `GLAD` to include prior information about the TFs if available. We assume that all the edges in the actual GRN have at least one gene which belongs to the list of TFs. We introduce a masking operation at every step of the unrolled iterations as shown in Algorithm 4), which eliminates all the unwanted edges that are between the non-TF nodes.

### B: Details of preprocessing data from BEELINE framework

<u>Preprocessing the real data:</u> For each gene expression data and its corresponding network, we do the following preprocessing to prepare our data. We first sort all the genes according to their variance and select the top 500 varying genes. We also have access to a list of known TFs. We only consider all the TFs whose variance had p-value at most 0.01. Now, we find the intersection between the top 500 varying genes and all the TFs to find the subset of genes which act as the TF, Table 3 in Supplementary-B. We then select the sub-graph of top 500 varying genes from the underlying GRN as our ground truth for evaluation.

**Algorithm 4:**
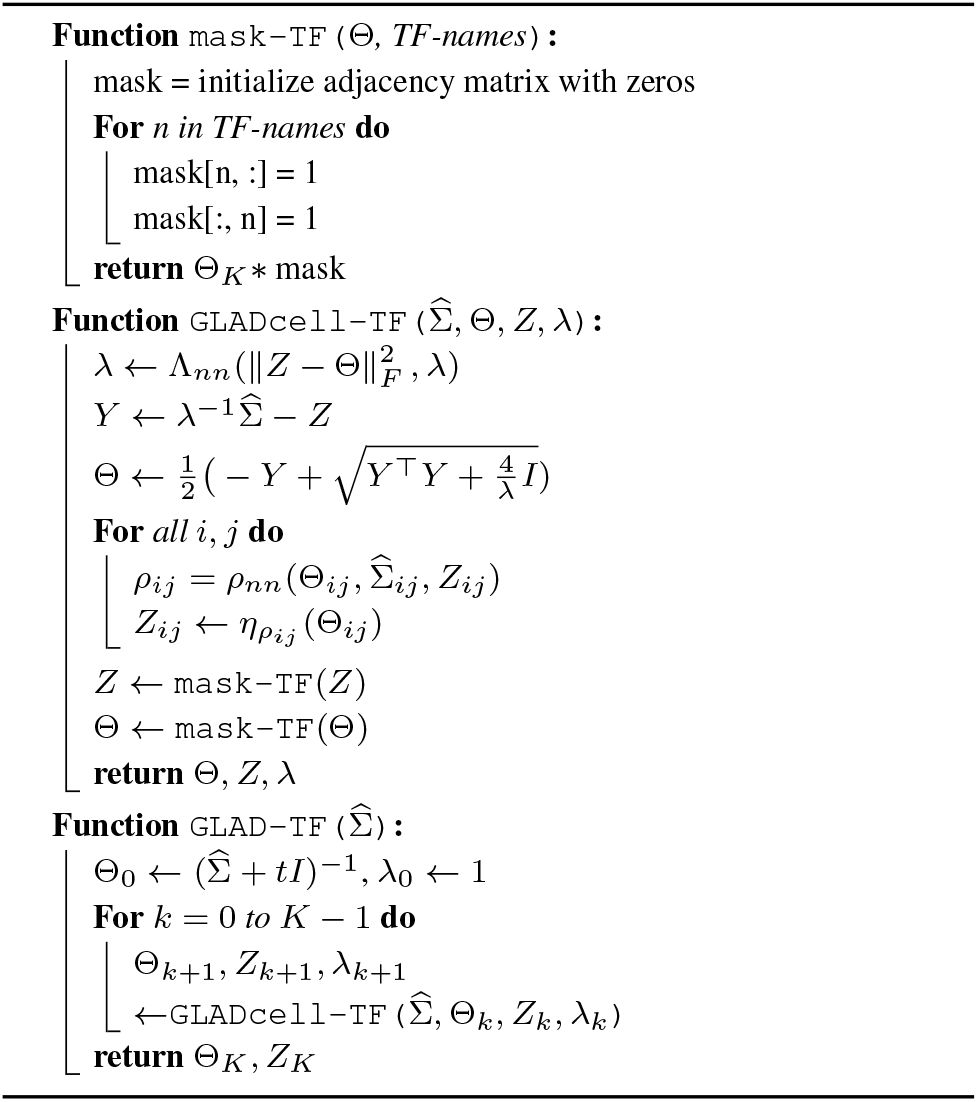
GLAD-TF

**Table 3.** Details of expression data from the BEELINE framework. To total number of genes for each data is 500 (highest varying genes

| Data | # TFs | # Expts |
| --- | --- | --- |
| mDC | 42 | 383 |
| mESC | 100 | 421 |
| mHSC-E | 69 | 1071 |
| mHSC-GM | 74 | 889 |
| mHSC-L | 83 | 847 |
| hESC | 86 | 758 |
| hHep | 56 | 425 |

